# Targeted protein degradation using degradFP in *Trypanosoma brucei*

**DOI:** 10.1101/2022.06.01.494430

**Authors:** Midori Ishii, Bungo Akiyoshi

## Abstract

Targeted protein degradation is an invaluable tool in studying the function of proteins. Such a tool was not available in *Trypanosoma brucei*, an evolutionarily divergent eukaryote that causes human African trypanosomiasis. Here, we have adapted degradFP (degrade Green Fluorescent Protein), a protein degradation system based on the SCF E3 ubiquitin ligase complex and anti-GFP nanobody, in *T. brucei*. As a proof of principle, we targeted a kinetoplastid kinetochore protein (KKT3) that constitutively localizes at kinetochores in the nucleus. Induction of degradFP in a cell line that had both alleles of KKT3 tagged with YFP caused more severe growth defect than RNAi in procyclic (insect form) cells. degradFP also worked on a cytoplasmic protein (COPII subunit, SEC31). Given the easiness in making GFP fusion cell lines in *T. brucei*, degradFP can serve as a powerful tool to rapidly deplete proteins of interest, especially those with low turnover rates.

## Introduction

Kinetoplastids are a group of unicellular flagellated eukaryotes found in diverse environmental conditions (d’Avila-Levy et al., 2015). They belong to the phylum Euglenozoa (Discoba/Excavata) and are evolutionarily distant from commonly studied model eukaryotes such as yeasts, worms, flies, and humans (Opisthokonta) (Keeling and Burki, 2019). Furthermore, it has been proposed that kinetoplastids may represent one of the earliest-branching eukaryotes based on a number of unique molecular features (Cavalier-Smith, 2010). Understanding their biology could therefore provide insights into the extent of conservation/divergence among eukaryotes and lead to a deeper understanding of biological systems and evolution of eukaryotes. Importantly, three neglected tropical diseases are caused by parasitic kinetoplastids: African trypanosomiasis, Chagas disease, and leishmaniasis (Rao et al., 2019; Horn, 2022). Human African trypanosomiasis (sleeping sickness) is caused by *Trypanosoma brucei*, which also causes the cattle disease, nagana, that leads to weight loss and anemia in livestock and imposes a huge burden on economic development in affected regions. Understanding the biology of kinetoplastids could facilitate the design of new drugs against kinetoplastid parasites.

Inducible depletion of a target protein is an essential tool in biology (Prozzillo et al., 2020). In *Trypanosoma brucei*, this can be achieved by RNAi (Ngô et al., 1998; Alsford et al., 2011) and Tet-off system (Merritt and Stuart, 2013) at the RNA level, as well as by conditional knockout at the gene level using Cre-LoxP (Kim et al., 2013) and CRISPR/Cas9 (Beneke et al., 2017). Although powerful in many cases, these approaches are not efficient in reducing the level of proteins that have slow turnover rates. Targeted degradation tools could circumvent this problem and have been used in other organisms (Uhlmann et al., 2000; Nishimura et al., 2009; Madeira da Silva et al., 2009; Damerow et al., 2015; Wheeler et al., 2015; Nabet et al., 2018). However, such tools were not available in *Trypanosoma brucei*, to our knowledge.

In this study, we have adapted the degradFP (degrade Green Fluorescent Protein) system which was originally established in *Drosophila melanogaster* (Caussinus et al., 2011). It relies on the expression of VhhGFP4 fused with a truncated F-box protein. VhhGFP4 is an anti-GFP nanobody that recognizes GFP and some derivatives such as YFP and Venus (Saerens et al., 2005), while an F-box protein is a substrate-recognition subunit of the SCF E3 ubiquitin ligase complex that catalyzes the ubiquitylation of target proteins (Petroski and Deshaies, 2005). In degradFP, a substrate-recognition domain of an F-box protein is replaced by VhhGFP4 so that GFP-fusion proteins are ubiquitylated by the SCF complex, leading to their degradation via the 26S proteasome pathway (Caussinus and Affolter, 2016). degradFP or modified versions have been used in mammalian cells, *C. elegans*, zebrafish, and plants (Shin et al., 2015; Wang et al., 2017; Yamaguchi et al., 2019; Sorge et al., 2021). Here, we show that degradFP successfully depletes a kinetochore protein and a COPII subunit in the procyclic form of *T. brucei* cells.

## Methods

We expressed the first 200 amino acids of Tb927.5.700 from *Trypanosoma brucei* (named FBP75 herein for <u>F</u>-<u>b</u>ox <u>p</u>rotein <u>75</u> kDa) that contain a putative F-box fused with the anti-GFP nanobody VhhGFP4 (Saerens et al., 2005) (Figure 1A, B). We made two constructs: one with a nuclear localization signal (NLS) to target nuclear proteins (pBA2675), and one without an NLS to target cytoplasmic proteins (pBA2705). In each case, the fusion protein was expressed from a derivative of pDEX777 that integrates at the 177 bp locus and allows doxycycline-inducible expression (Kelly et al., 2007; Nerusheva and Akiyoshi, 2016). Induction of these degradFP systems in wild-type procyclic cells with 1 µg/mL doxycycline did not cause any growth defect (Figure 1C).

**Figure 1.**
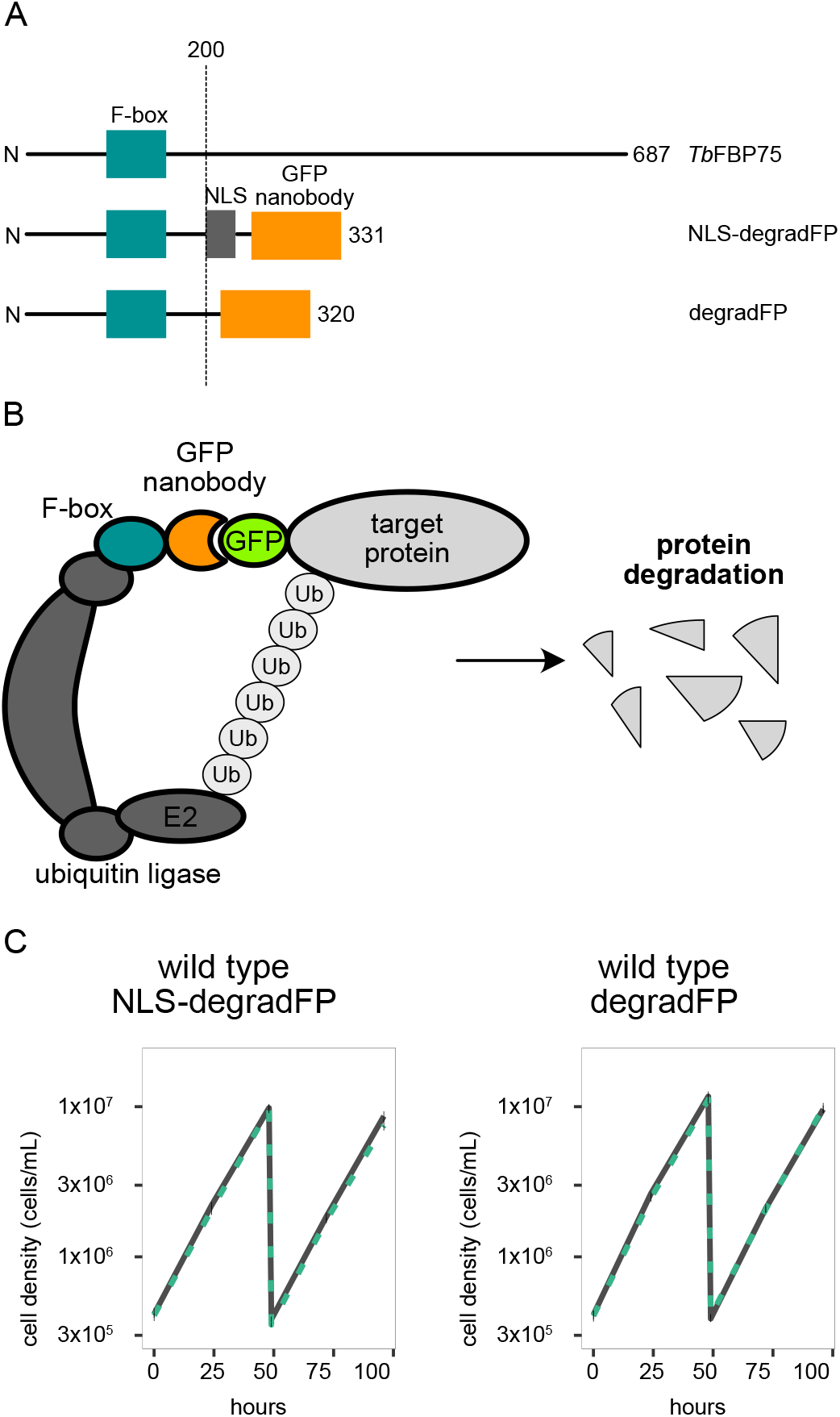
degradFP in *Trypanosoma brucei*. (A) Schematic of *Tb*FBP75, NLS-degradFP, and degradFP, highlighting the putative F-box domain, NLS, and GFP nanobody (vhhGFP4). (B) degradFP forms a complex with an endogenous ubiquitin ligase complex which transfers the ubiquitin to the target protein tagged with GFP. Ubiquitin-tagged target proteins are then degraded by the 26S proteasome. (C) Growth curve for wild-type procyclic cells with NLS-degradFP (left) or degradFP (right). degradFP was induced with 1 μg/ml doxycycline and cultures were diluted at day 2. Gray lines are uninduced controls. Green dushed lines are doxycycline-treated cells. N=3. Error bars are SEM. Cell line: BAP2395, BAP2511

## Results

We first targeted KKT3, a kinetochore protein that constitutively localizes at kinetochores (Akiyoshi and Gull, 2014). KKT3 does not show any obvious fluctuation in its abundance during the cell cycle, implying that it is a stable protein. In fact, the half-life of KKT3 has been estimated to be much longer than transiently-localized kinetochore proteins (Tinti et al., 2019). Both alleles of KKT3 were C-terminally tagged with YFP using a PCR-based method in one transfection step (Dean et al., 2015). We found that induction of degradFP in this cell line caused more severe growth defects than RNAi (Figure 2A and B) (Marcianò et al., 2021). Microscopy analysis confirmed significant depletion of KKT3 at 6 hours (Figure 2B). The fact that induction of degradFP in wild-type cells did not cause any growth defect (Figure 1C) means that the observed growth defect was due to specific degradation of YFP-tagged KKT3.

**Figure 2.**
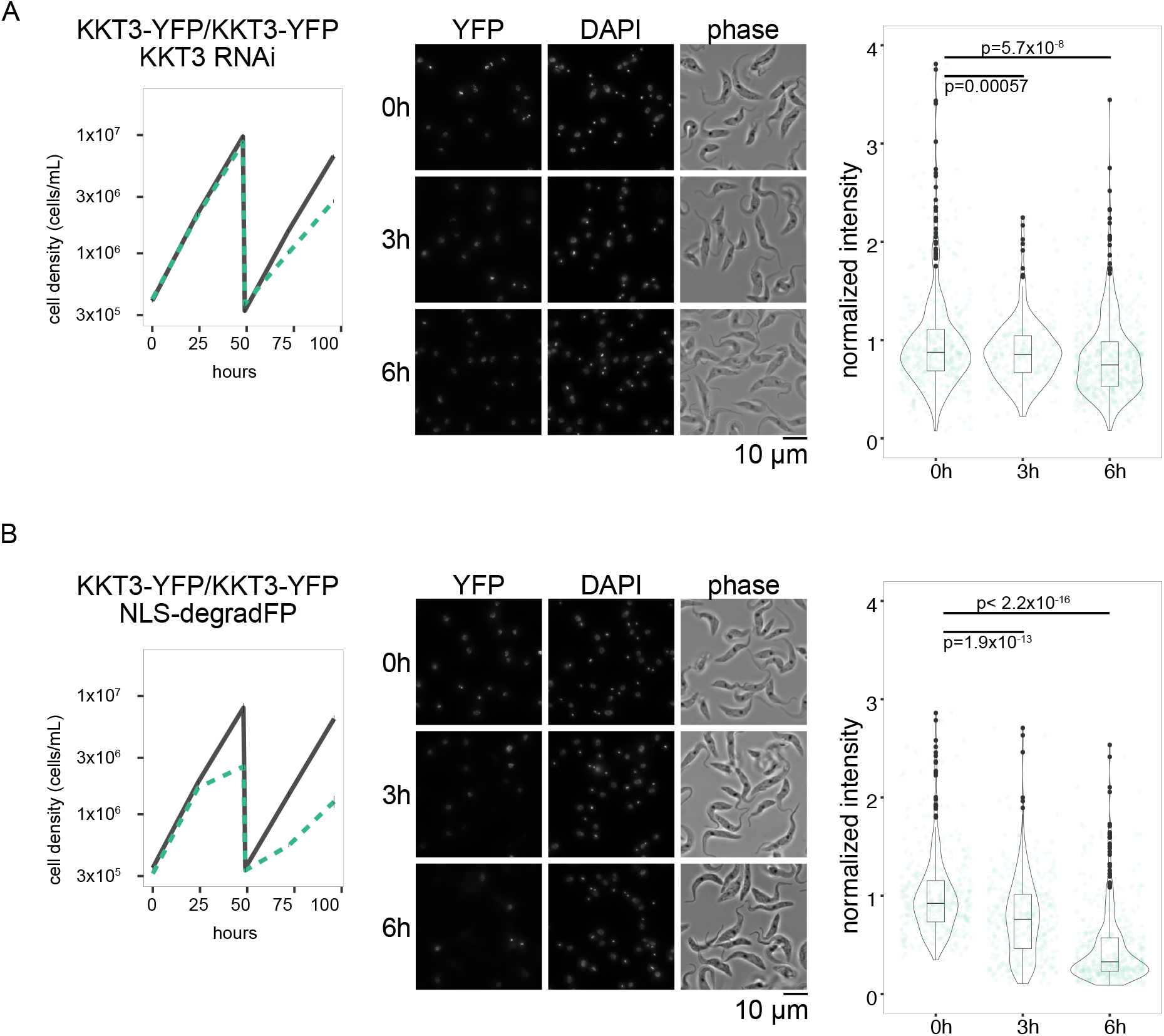
degradFP depletes a nuclear protein KKT3 more efficiently than RNAi. (A) KKT3 knockdown by RNAi. Cell line: BAP2512, (B) KKT3-YFP depletion by degradFP with NLS. Cell line: BAP2513. (Left) Growth curve. RNAi or degradFP was induced with 1 μg/ml doxycycline and cultures were diluted at day 2. Gray lines indicate uninduced controls. Green dushed lines are doxycycline-treated cells. N=4 (RNAi) and 3 (degradFP). Error bars are SEM. (Centre) Examples of cells at 0 h, 3 h, and 6 h after induction. YFP and DAPI images are maximum intensity projection. (Right) Plot of total YFP signal inside the nucleus (>239 cells in each condition). Data was normalized with the mean value at 0 h. P-values were calculated by Welch two sample t-test.

We next targeted a cytoplasmic protein SEC31 using a degradFP construct that lacks an NLS. SEC31 is a subunit of COPII and localizes at the ER exit site (Hu et al., 2016). Both alleles of SEC31 were C-terminally tagged in a CRISPR cell line (Beneke et al., 2017). Induction of degradFP caused a strong growth defect (Figure 3), which is apparently more severe than RNAi-mediated depletion of SEC31 reported in a previous study (Hu et al., 2016). These results therefore show that degradFP can efficiently deplete both nuclear and cytoplasmic proteins in *T. brucei*.

**Figure 3.**
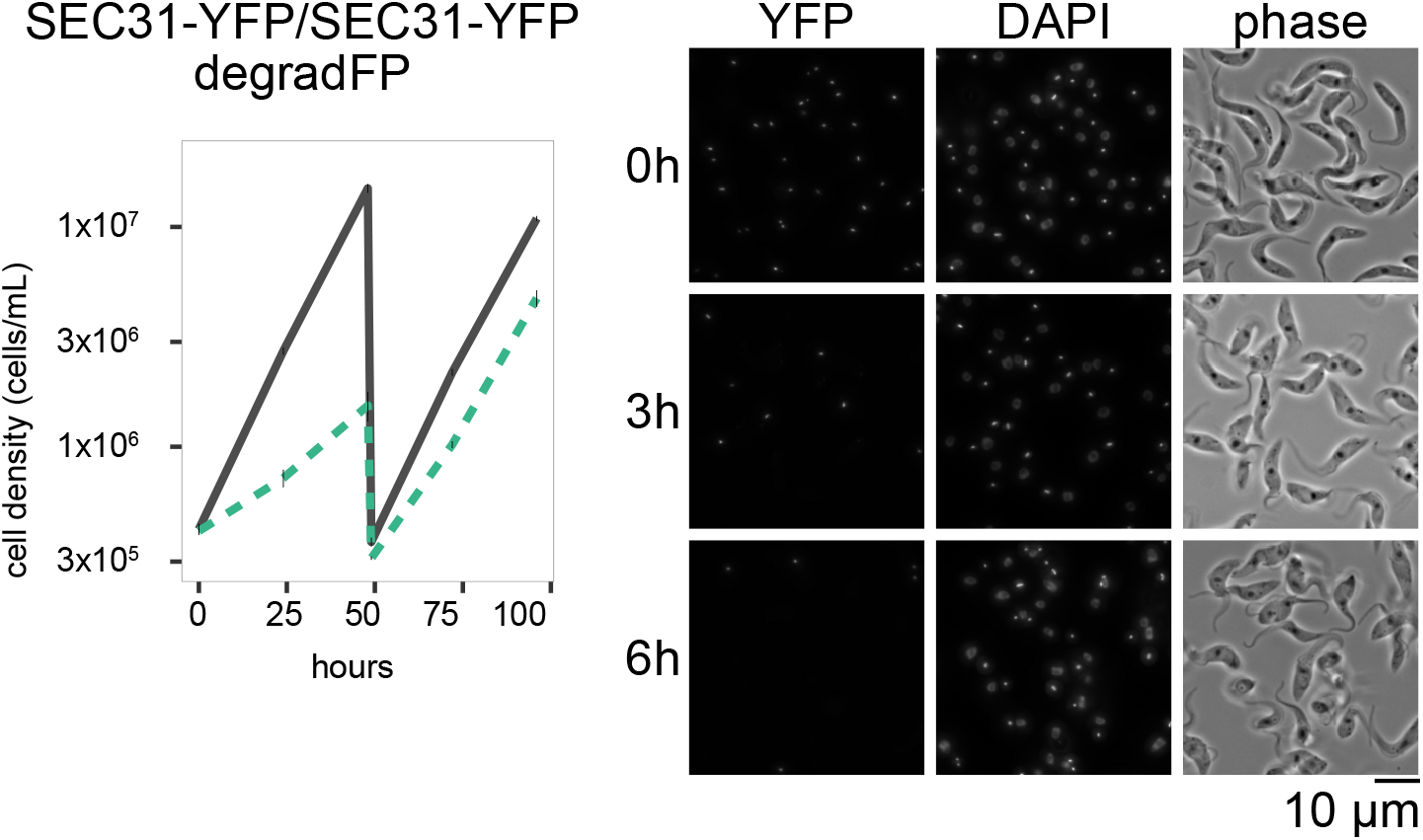
Depletion of a cytoplasmic protein SEC31 by degradFP. (Left) Growth curve. degradFP was induced with 1 μg/ml doxycycline and cultures were diluted at day 2. Gray line indicates an uninduced control. Green dushed line is doxycycline-treated cells. N=4. Error bars are SEM. (Right) Examples of cells at 0 h, 3 h, and 6 h after induction. YFP and DAPI images are maximum intensity projection. Cell line: BAP2518.

## Discussion

In *T. brucei*, it is easy to tag genes at the endogenous locus using plasmid- or PCR-based methods (Kelly et al., 2007; Dean et al., 2015; Beneke et al., 2017; Kovářová et al., 2022). Taking advantage of the inducible expression system (Wirtz and Clayton, 1995; Poon et al., 2012), we have shown that degradFP can induce targeted protein degradation in *T. brucei*. The depletion kinetics is faster than the RNAi-mediated depletion method at least for KKT3. Our results therefore show that degradFP can be a powerful tool in characterizing depletion phenotypes in *T. brucei*. It is, however, important to note that degradFP has some limitations. For example, it has been suggested that degradFP does not work if GFP is not accessible (Caussinus et al., 2011; Caussinus and Affolter, 2016). Furthermore, it is essential that GFP-fusion proteins retain enough functionality to support cell growth because the degradFP system utilizes the VhhGFP4 nanobody that recognizes GFP or its derivatives. If necessary, this system could be modified to use nanobodies against other epitope tags or even the protein of interest itself to induce degradation of the target.

Function of the F-box protein used in this study (FBP75) remains unknown. We also do not know which SKP1 or cullin proteins interact with FBP75 and whether those proteins are expressed in other life stages. It therefore remains unknown whether FBP75-based degradFP works in bloodstream form cells. If it does not work, other F-box proteins could be utilized to deplete proteins of interest in bloodstream form cells (Benz and Clayton, 2007; Rojas et al., 2017). In any case, it is our hope that degradFP will prove to be a useful protein degradation tool to facilitate studies of *Trypanosoma brucei*.

## Materials and methods

### Plasmids

All plasmids used in this study are listed in Table 1. To make pBA2675 (Inducible expression of FBP75^1−200^-NLS-VhhGFP4: NLS-degradFP for nuclear proteins), synthetic DNA BAG181 (GeneArt, Thermo Fisher) was digested with HindIII/BamHI and subcloned into pBA310 cut with the same enzymes. The NLS sequence was derived from the La protein (Marchetti et al., 2000). To make pBA2705 (Inducible expression of FBP75^1−200^-VhhGFP4: degradFP for cytoplasmic proteins), NLS was removed from pBA2675 by PCR with BA3647/BA3648. To make pBA1061 (hairpin RNAi targeting 2562–3072 bp of the KKT3 coding sequence), synthetic DNA BAG55 (GeneArt, Thermo Fisher) was digested with HindIII/BamHI and subcloned into pBA310 cut with the same enzymes.

**Table 1.** Plasmids used in this study

| Name | Description |
| --- | --- |
| pPOTv7 (eYFP, Hyg) | PCR-based eYFP-tagging vector, hygromycin marker (Dean et al., 2015) |
| pPOTv7 (eYFP, G418) | PCR-based eYFP-tagging vector, hygromycin marker (Dean et al., 2015) |
| pBA310 | Inducible expression vector, integrate at 177 bp, phleomycin marker (Nerusheva and Akiyoshi, 2016) |
| pBA1061 | Inducible expression of KKT3 hairpin RNAi (targeting 2562–3072 bp of KKT3 coding sequence), integrate at 177 bp, phleomycin marker |
| pBA2675 | Inducible expression of FBP75<sup>1–200</sup>-NLS-VhhGFP4, integrate at 177 bp, phleomycin marker (degradFP for nuclear proteins) |
| pBA2705 | Inducible expression of FBP75<sup>1–200</sup>-VhhGFP4, integrate at 177 bp, phleomycin marker (degradFP for cytoplasmic proteins) |

### Trypanosome cells

All cell lines used in this study were derived from the TREU 927 procyclic form cells and are listed in Table 2. SmOxP927 expresses Tet repressor and T7 RNA polymerase (Poon et al., 2012), while PCF 1339 expresses Tet repressor, T7 RNA polymerase, and the Cas9 nuclease (Beneke et al., 2017). Cells were grown at 28°C in SDM-79 medium (Life Technologies, Thermo Fisher) supplemented with 10% heat-inactivated fetal calf serum (Sigma) and 7.5 µg/mL hemin, as well as puromycin (Sigma) and appropriate drugs (Brun and Schönenberger, 1979).

**Table 2.**
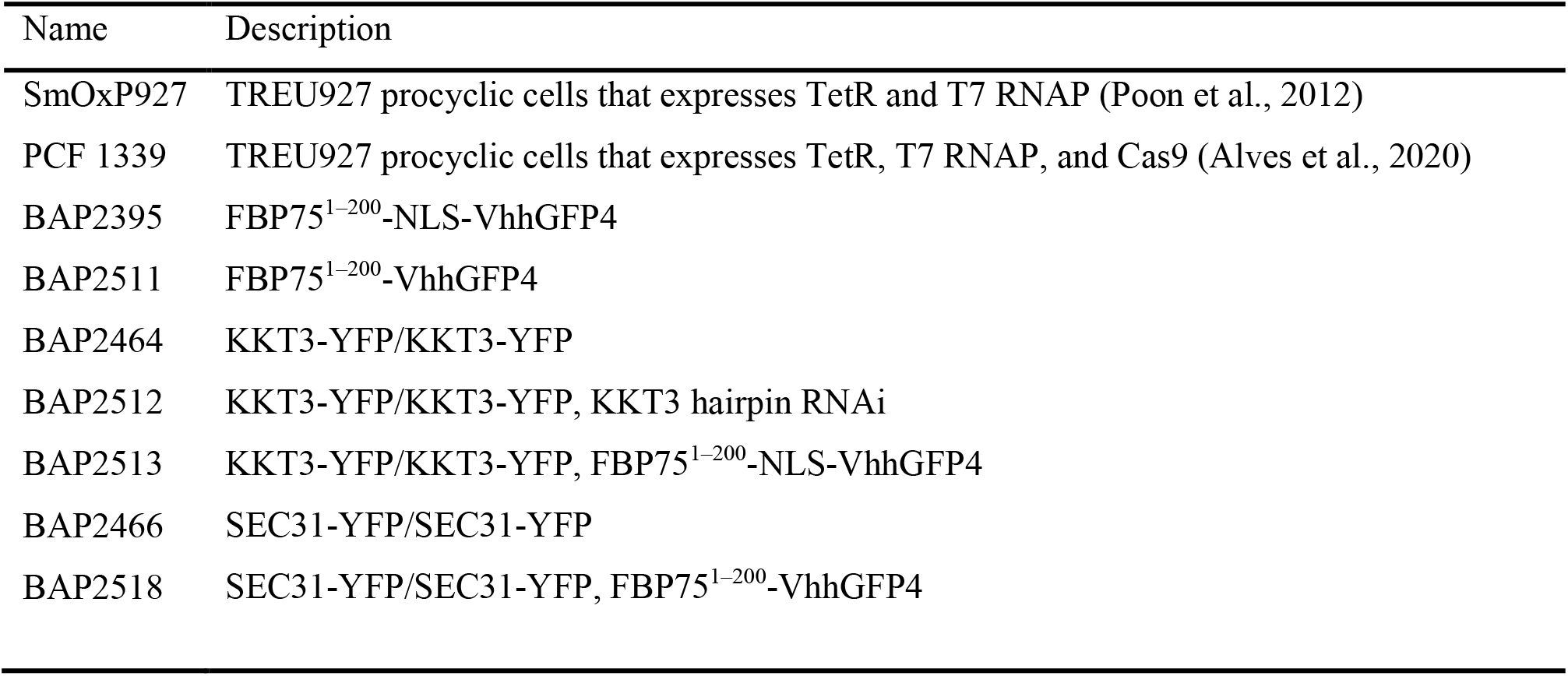
Trypanosome cell lines used in this study

**Table 3.**
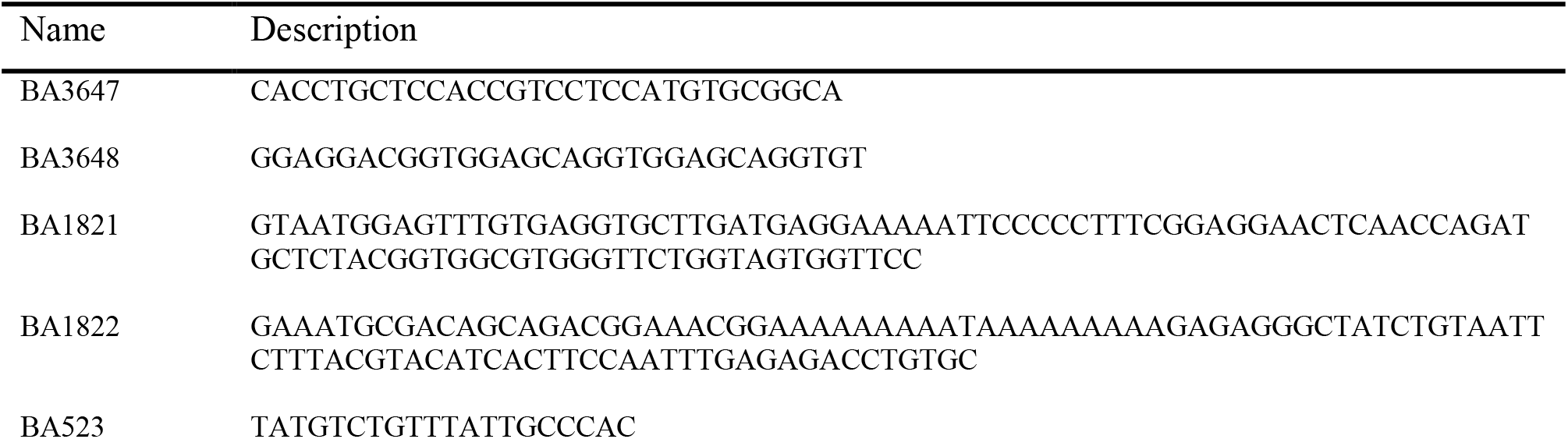
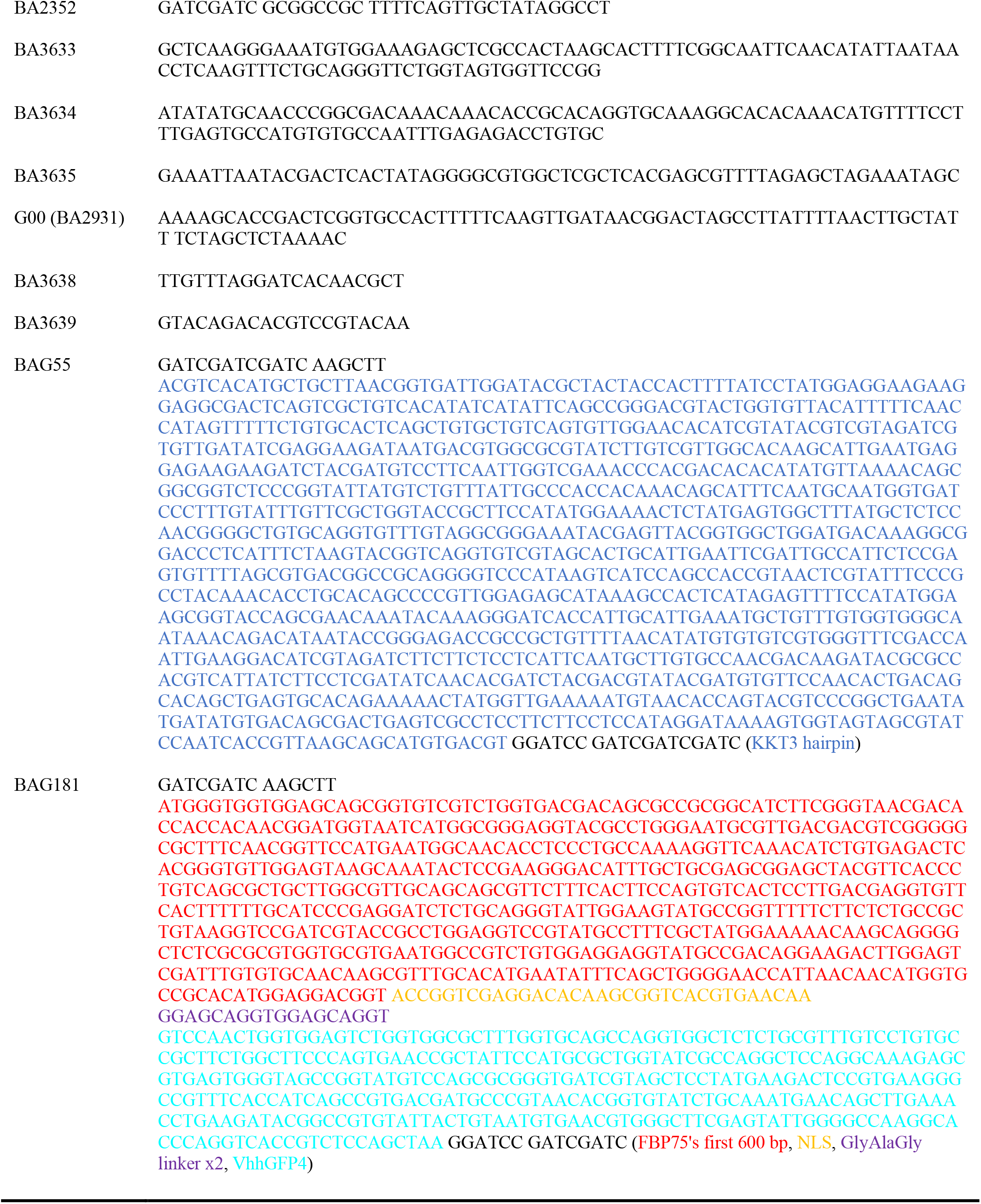
Sequence of primers and synthetic DNA used in this study

To make the homozygous KKT3-YFP cell line, two YFP-tagging cassettes were amplified from pPOTv7 (YFP, Hyg) or pPOTv7 (YFP, G418) (Dean et al., 2015) by PCR using BA1821/BA1822 and transfected into SmOxP927 (Poon et al., 2012) by electroporation. Transfected cells were selected by addition of 50 μg/mL hygromycin (Sigma) and 30 μg/mL G418 (Sigma) and cloned by dispensing dilutions into 96-well plates. Clones were screened by diagnostic PCR of genomic DNA using BA523/BA2352.

To make the homozygous SEC31-YFP cell line, donor DNA templates amplified from pPOTv7 (YFP, Hyg) or pPOTv7 (YFP, G418) (Dean et al., 2015) with BA3633/BA3634 and sgRNA template amplified with BA3635/G00 were co-transfected into PCF 1339 (Beneke et al., 2017; Alves et al., 2020) by electroporation. Transfected cells were selected by addition of 50 μg/mL hygromycin (Sigma) and 30 μg/mL G418 (Sigma) and cloned by dispensing dilutions into 96-well plates. Clones were screened by diagnostic PCR of genomic DNA using BA3638/BA3639. RNAi and degradFP constructs were linearized by NotI and transfected into YFP-tagged cell lines or SmOxP927 by electroporation and selected by addition of 5 µg/mL phleomycin (Sigma). For induction of degradFP or RNAi, doxycycline (Sigma) was added to the medium to a final concentration of 1 μg/mL. Cell growth was monitored using a CASY cell counter (Roche) and plotted with ggplot in R.

### Fluorescence microscopy

Cells were harvested by centrifugation at 800 g for 5 min, washed in PBS, settled onto glass slides for 5 min, and fixed with 4% paraformaldehyde for 5 min. Following three washes in PBS (5 min each), cells were permeabilized with 0.1% NP-40 in PBS for 5 min, washed three times in PBS (5 min each), and embedded in mounting media (1% 1,4-Diazabicyclo [2.2.2]octane (DABCO), 90% glycerol, and 50 mM sodium phosphate pH 8.0) containing 100 ng/mL DAPI. Images were captured at room temperature on a Zeiss Axioimager.Z2 microscope (Zeiss) installed with ZEN using a Hamamatsu ORCA-Flash4.0 camera with 63x objective lenses (1.40 NA). 22 optical slices spaced 0.24 μm apart were collected. Images were processed in ImageJ/Fiji (Schneider et al., 2012). Maximum intensity projection images were generated by Fiji software (Schneider et al., 2012). Total YFP intensity in the nucleus was measured using 3D Objects Counter with default settings in Fiji as follows. DAPI images were first used to segment the nucleus by removing regions that have top 0.2% intensity (that correspond to kDNA signals) and then by selecting objects that have the size of nuclei (5.3–40 µm^3^). YFP total intensity inside the nucleus was measured using a redirect function in 3D Objects Counter. 3D Plots were made with ggplot in R.

## Competing Interests

We declare no competing interests.

## Acknowledgments

We thank Markus Affolter for advice. We also thank Jack Sunter, Sam Dean, and Tom Beneke for sharing reagents. pBA2675 (NLS-degradFP for nuclear proteins) and pBA2705 (degradFP for cytoplasmic proteins) are freely available upon request.

## Grant information

B. Akiyoshi was supported by a Wellcome Trust Senior Research Fellowship (grant 210622/Z/18/Z) and the European Molecular Biology Organization Young Investigator Program.

## Rights retention

This research was funded in whole or in part by Wellcome Trust (grant 210622/Z/18/Z). For the purpose of Open Access, the author has applied a CC BY public copyright licence to any Author Accepted Manuscript (AAM) version arising from this submission.

## Notes

### Competing Interest Statement

The authors have declared no competing interest.

